# microRNA seedless sites attenuate strong-seed-site-mediated target repression

**DOI:** 10.1101/837682

**Authors:** Xujun Wang, Jingru Tian, Peng Cui, Stephen Mastriano, Dingyao Zhang, Hongyu Zhao, Hui Lu, Ye Ding, Jun Lu

**Author notes:** These authors contributed equally to this work (Xujun Wang, Jingru Tian, Peng Cui, Stephen Mastriano). Correspondence should be addressed to: Jun Lu, 10 Amistad Street, Room 237C, New Haven, CT 06520.

## Abstract

MicroRNAs (miRNAs) regulate protein-coding gene expression primarily through cognitive binding sites in the 3’ untranslated regions (3′ UTRs). Seed sites are sequences in messenger RNAs (mRNAs) that form perfect Watson-Crick base-paring with a miRNA’s seed region, which can effectively reduce mRNA abundance and/or repress protein translation. Some seedless sites, which do no form perfect seed-pairing with a miRNA, can also lead to target repression, often with lower efficacy. Here we report the surprising finding that when seedless sites and seed sites are co-present in the same 3’UTR, seedless sites attenuate strong-seed-site-mediated target suppression, independent of 3′ UTR length. This attenuation effect is detectable in >70% of transcriptomic datasets examined, in which specific miRNAs are experimentally increased or decreased. The attenuation effect is confirmed by 3’UTR reporter assays and mediated through base-pairing between miRNA and seedless sites. Furthermore, this seedless-site-based attenuation effect could affect seed sites of the same miRNA or another miRNA, thus partially explaining the variability in target suppression and miRNA-mediated gene upregulation. Our findings reveal an unexpected principle of miRNA-mediated gene regulation, and could impact the understanding of many miRNA-regulated biological processes.

## Main Text

MicroRNAs (miRNAs) are ~ 22 nt small RNAs that regulate protein-coding gene expression primarily through cognitive binding sites in the 3’ untranslated regions (3′ UTRs). Principles that underlie efficient miRNA-mediated target repression are important in deciphering miRNA functions in diverse biological processes. However, existing rules fall short of fully explaining miRNA-mediated gene regulation. The seed site rule specifies that a miRNA can bind to target mRNA sequence through perfect Watson-Crick base-paring between miRNA’s seed region (nucleotides 2 to 7) and its target site, thus leading to target repression through miRNA-mediated recruitment of Argonaute (AGO) proteins and subsequent post-transcriptional target degradation and/or translational inhibition ^1,2^. Based on pairing patterns and nucleotide identity, seed sites are further classified into 8mer, 7mer-A1 and 7mer-M8 sites which elicit relatively efficient target downregulation (referred to collectively in this study as “strong seed sites”), and 6mer and offset 6mer sites which have generally weaker abilities to regulate target gene expression ^3^. In addition to seed sites, miRNA-target interaction could also occur through seedless sites (also known as non-canonical sites), in which the binding does not require perfect Watson-Crick pairing of the miRNA seed region ^4–13^. A fraction of seedless sites could also confer miRNA-mediated target repression, but the effectiveness of seedless site in suppressing target expression is often weaker and could depend on the location, sequence and structural context ^14–17^. Despite these progresses, it has also been well appreciated that overexpression or knockout of specific miRNAs will cause wide-spread changes in mRNA and protein levels, including both increased and decreased expression. Additionally, predicted target genes with the same type of strong seed sites are often regulated at different levels. Some targets were strongly repressed, some were weakly repressed, and others either were hardly repressed or even have enhanced expression. Such a wide spectrum of outcomes of miRNA-mediated target suppression hints at likely unknown principles that further govern miRNA-mediated target regulation.

We focused on the effects of seedless sites in 3’ UTRs, given that seedless sites are the least understood class of miRNA recognition sequences, and the significance of biochemically identified seedless sites has been dismissed by a recent report as having little regulatory effects ^16^. Specifically, we asked the question of when seedless sites coexist with seed sites on the same 3′ UTR, what could be the potential impact of seedless sites on seed-site mediated target repression (Figure 1A). To answer this question, we predicted both seed and seedless sites in 3′ UTRs across the human and mouse genomes based on our previously published methods ^18,19^ (see **Methods**). We then analyzed the association between predicted miRNA recognition sites and transcriptomic changes from published datasets in which specific miRNAs were overexpressed, knocked out (KO) or knocked down (KD). Of note, it has been reported that RNA-level changes account for the majority of miRNA-mediated effects ^20^. Consistent with target repression by strong seed sites, when human miR-124 was overexpressed in HeLa cells, predicted strong targets of miR-124-3p (with 3’UTRs containing 8mer, 7mer-A1 or 7mer-M8 site(s)) were significantly down-regulated compared to “non-seed genes” (defined in this manuscript as those 3′ UTRs without any types of seed sites) (Figure 1B). Previous studies examining two or more seed sites on the same 3′ UTR have found frequent co-operation between the seed sites, leading to additive or synergistic miRNA-mediated target suppression (e.g. ^21–25^). Indeed, we observed that miRNAs with two or more strong seed sites in the 3′ UTR were significantly more strongly suppressed by miRNA expression than those with single strong seed sites (**Supplementary Figure S1A**). Based on the co-operative logic, one would expect that the presence of seedless sites in the same 3′ UTR with strong seed sites could either lead to enhanced cooperative suppression or have the same levels of suppression as those regulated by the strong-seed sites alone. Surprisingly, however, we observed that strong-seed targets with more seedless sites (top 1/3 of predicted seedless site counts) showed significantly less miRNA-mediated suppression than those with fewer seedless sites (bottom 1/3) (Figure 1B), with a graded response dependent on the number of predicted seedless sites (**Supplementary Figure S1B**). Similar effects were seen when miR-122 knockout murine liver samples were compared to wild-type liver, with reduced de-repression of strong-seed-targets in the presence of more predicted seedless sites (Figure 1C, **Supplementary Figures S1C & 1D**). On average, target repression was weakened by 45.6% for miR-124 over-expression and by 48.7% for miR-122 knockout, when comparing strong-seed targets with top 1/3 seedless sites to those with bottom 1/3 seedless sites (Figure 1D, 1E). Since these unexpected results can be potentially explained by a model in which the presence of seedless sites attenuates the strong-site-mediated target repression, we thus refer to such effects below as “attenuation effects” for simplicity. Of note, the above attenuation effects cannot be explained by any opportunistic association, or the lack thereof, between the number of predicted seedless sites with strong seed site types (i.e. 8mer, 7mer-m8 and 7mer-A1) or strong site count (**Supplementary Figure S1E, 1F**). To more comprehensively quantify the attenuation effect by the number of predicted seedless sites, we performed linear regression modeling across 21 randomly collected public datasets on 18 miRNAs. For each miRNA overexpressed or knocked down/knocked out, we used single-variable linear regression to quantify the per count influence of seedless sites on strong-site-mediated target repression. We observed significant attenuation effects in 15 out of 21 datasets (including 4 out of 5 miRNA KD/KO datasets) (**Supplementary Table 1**). For the remaining six datasets, no significant association was observed for either attenuation or enhancement by seedless site count on strong-seed-site-mediated target suppression. Taken together, the data above reveal significant associations between the number of predicted seedless sites and the attenuation of strong-site-mediated target repression for the majority of miRNAs examined.

**Figure 1.**
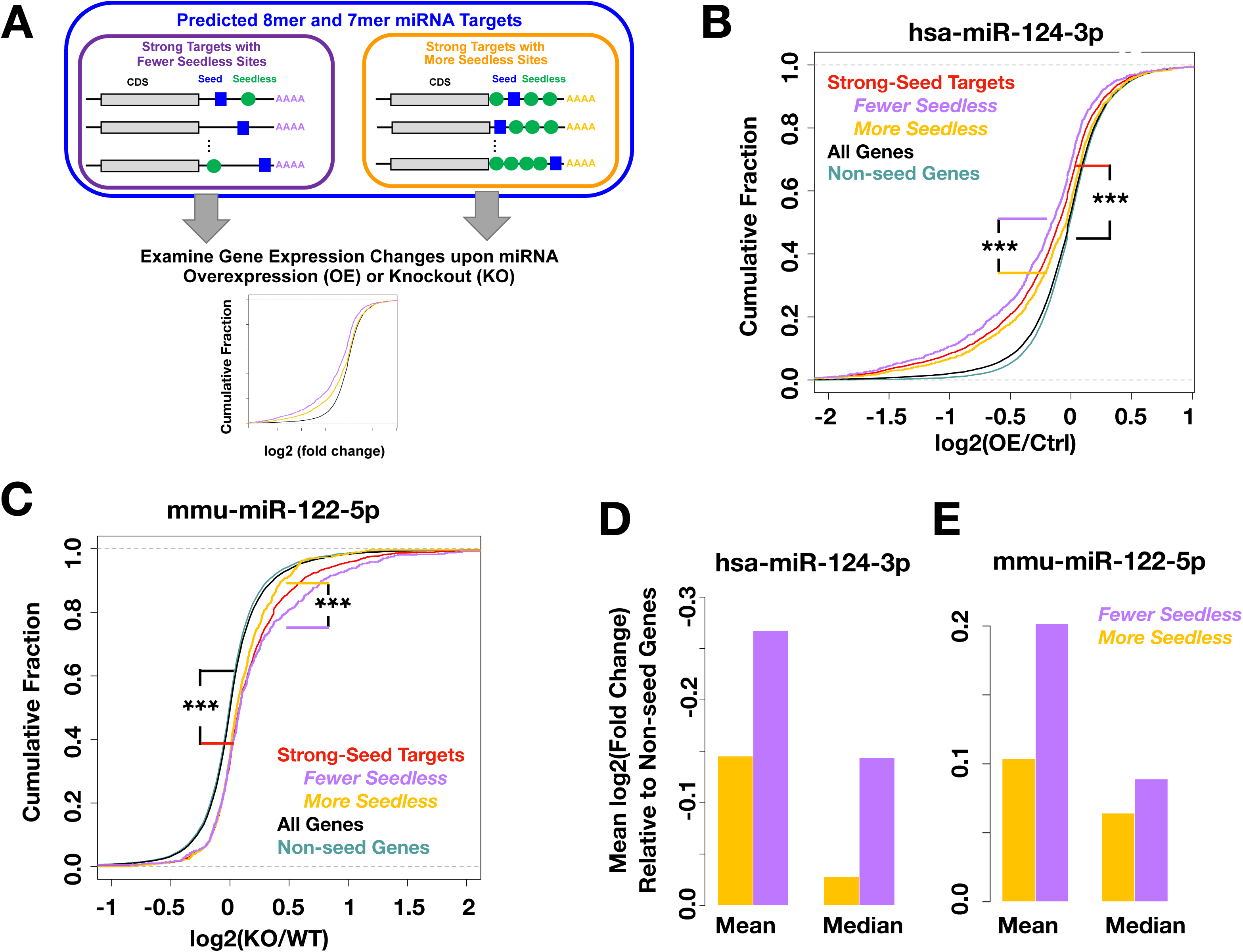
Higher miRNA seedless site count is associated with an attenuation of strong-seed-site-mediated target suppression. **(A)** A schematic of analysis, in which genes with predicted strong seed sites (8mer and 7mer sites) in the 3’UTR were further divided into those with more and fewer predicted seedless sites (top and bottom 1/3 of genes, respectively), and compared for miRNA-induced gene expression changes. **(B, C)** Cumulative distribution function plots for the indicated groups of genes comparing gene expression in (B) miR-124 overexpression (OE) versus control (Ctrl) HeLa cells, and (C) miR-122 knockout (KO) liver versus wildtype (WT) liver. Non-seed genes refer to genes without predicted seed sites in the 3’UTR. Legends are color-coded to match the line colors. ***: p<0.001. **(D, E)** miRNA-induced gene expression changes were plotted for genes with predicted strong seed sites containing more or fewer seedless sites for data in (B) and (C) respectively. Expression changes were calculated by comparison to non-seed genes, and both mean and median expression changes were plotted.

It has been reported that longer 3′ UTRs are associated with less experimentally observable miRNA binding ^26–28^, without clear mechanisms, whereas a positive association between 3′ UTR length and the level of miRNA-mediated target suppression has also been described ^28,29^. Indeed, 3′ UTR length has been incorporated as a feature in several computational models for miRNA-mediated target suppression ^9,16,30^. Since we often observed a positive correlation between the number of predicted seedless sites and 3′ UTR length, for example for hsa-miR-124 and mmu-miR-122 (**Supplemental Figure S2A, 2B**), we asked whether the attenuation effect by the number of seedless sites is merely secondary to a primary effect of 3′ UTR length, or instead is independent from any effect of the 3′ UTR length. To address this question, we first designed an analysis in which the length of the 3′ UTR was fixed but the number of seedless sites were variable. We have previously published data from 3′ UTR reporter assays which quantified the regulatory effects of 460 individually overexpressed miRNA constructs on human and mouse Tet2 3′ UTR reporters ^31^. This dataset has the added advantage of isolating the miRNA-mediated regulatory effect on the 3′ UTR from miRNA-induced secondary effects such as those through transcription. In examination of the regulation of the human TET2 3′ UTR, which is of the same 3′ UTR length across all assayed miRNAs, we subcategorized miRNAs that have predicted strong target sites in the TET2 3′ UTR into those with more seedless sites (top 1/3) or those with fewer (bottom 1/3). MiRNAs with more seedless-sites showed significantly reduced 3′ UTR repression than those with fewer seedless-sites (Figure 2A). Similar findings were observed for mouse Tet2 3′ UTR (Figure 2B), despite that the lists of miRNAs regulating human and mouse Tet2 3′ UTRs were non-overlapping ^31^. These data indicate that the attenuation effects by seedless sites is independent of 3′ UTR length.

**Figure 2.**
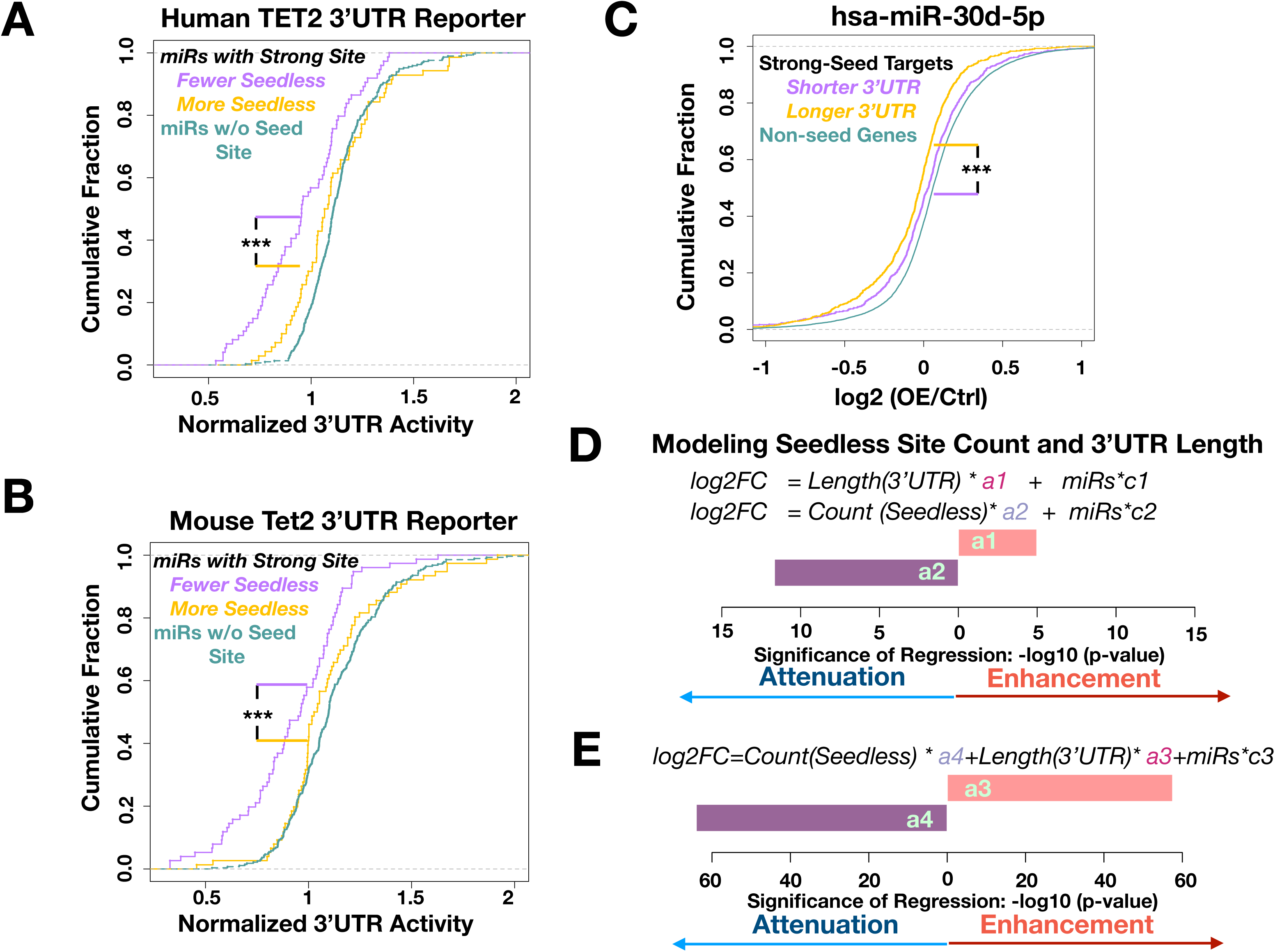
Seedless-site-based attenuation effect is independent from 3’UTR length. **(A, B)** A dataset in which the effects of ~460 miRNAs on (A) human TET2 3’UTR reporter and (B) mouse Tet2 3’UTR reporter, both of which have fixed 3’UTR lengths, was analyzed. Normalized 3’UTR reporter activities, with numbers below 1 reflecting a suppressing effect and numbers above 1 for an enhancing effect, were examined for miRNAs that have predicted strong seed sites in the 3’UTR or those without seed sites. Those with seed sites were further divided into those having more or fewer predicted seedless sites for the same miRNA (top and bottom 1/3 of miRNAs, respectively). ***: p<0.001. **(C)** An example of longer 3’UTR associated with stronger miRNA-seed-site-mediated target suppression. A cumulative distribution function plot for the indicated groups of genes comparing gene expression in miR-30d overexpression (OE) versus control (Ctrl) LNCaP cells. p<0.001. **(D, E)** Linear regression modeling of the effects of seedless site count and 3’UTR length on strong-seed-site-mediated target suppression. (D) Two separate linear regression models (indicated at the top) were applied to a meta-analysis on 21 miRNA perturbation datasets. The direction of association (enhancing or attenuating strong-seed-site-mediated target suppression) is indicated at the bottom. The lengths of the bars reflect the significance of association. (E) A combined linear regression model involving both seedless site count and 3’UTR length was analyzed similar to (D).

To further examine the relationship between seedless sites and 3′ UTR lengths on miRNA-mediated gene expression, we performed single-variable linear regression to model the effect of 3’UTR length on strong-site-mediated target repression in the same 21 datasets analyzed above. Results show that while 3′ UTR length was negatively associated with strong-seed-site-mediated target suppression in 12 out of 21 datasets (**Supplementary Table 2, Supplementary Figure S2C, 2D**), an opposite effect in which longer 3′ UTRs are associated with stronger target repression could be observed for 4 out of the 21 datasets (**Supplementary Table 2**). For instance, when miR-30d was overexpressed in the prostate DU145 cell line, strong targets with longer 3’UTRs were significantly more downregulated (Figure 2C). We then performed a meta-analysis to include all 21 miRNA datasets in a multi-variable linear regression model, using the 3′ UTR length and miRNA identity as variables. In this meta-analysis, a negative association between 3′ UTR length and strong-seed-site-mediated gene suppression could not be observed (Figure 2D). In contrast, the same multi-variable regression analysis for the number of seedless sites revealed a significant attenuation effect (Figure 2D). These data indicate that the influence of 3’UTR length is not as robust as the attenuation effects by seedless sites. To examine this notion further, we modeled both 3′ UTR length and the number of seedless sites in a multi-variable linear regression and observed a significant attenuation effect by the number of seedless sites, but failed to observe a similar effect by the 3′ UTR length (Figure 2D). These data further support that the attenuation effect by the number of seedless sites is independent from any effect of 3′ UTR length.

Could the attenuation effect be observed not only with predicted seedless sites, which may have high false-positive levels, but also with experimentally validated seedless binding sites for miRNAs? To answer this question, we focused on a published dataset in which miRNA-binding sites (a.k.a. miRNA response elements or MREs) were experimentally mapped through transfecting and pulling down biotinylated hsa-miR-522-3p in breast cancer MDA-MB-468 cells ^4^. The choice of this dataset instead of CLIP data with AGO immunoprecipitation (IP) was because of the confidence of the annotated seedless sites originating from the transfected miRNA, and because several reports have suggested the existence of an AGO-independent pool of miRNAs in mammalian cells ^32–34^ and thus procedures involving AGO IP might have missed binding by such a fraction of miRNAs. We first set out to better understand the relationship between MRE-verified targets and our target predictions. Among 2,820 predicted strong-seed targets, 551 showed predicted sites overlapping the MRE collection (Figure 3A). These overlapping targets were significantly downregulated upon ectopic miR-522 expression as expected (Figure 3B). However, 2,269 of the predicted strong-seed targets did not have overlapping MREs, but showed significant downregulation, albeit with a reduced mean repression (~43%) compared to the overlapping targets (Figure 3B, 3C). These data strongly suggest that the biochemically detectable miRNA binding site collection had a substantial level of false negatives and possibly have missed many functional but weak or transient binding events. Among 891,698 predicted seedless sites, 1,575 overlapped with the MRE collection (Figure 3D). The level of detectable miRNA occupancy for predicted seedless sites was substantially lower than that for strong-seed sites (0.18% vs 19.5%), suggesting either the existence of a high level of false positives in seedless site prediction or that miRNA binding to most seedless sites are transient and weak thus evading biochemical detection. Nevertheless, 3′ UTRs with higher numbers of predicted seedless sites had a higher probability of observing MREs (Figure 3E), suggesting that our use of the number of predicted seedless sites (in evaluating attenuation effects above) could reflect the likelihood of miRNA binding to seedless sites. We next asked whether we could observe an attenuation effect of seedless sites on strong-seed-site mediated target repression using experimentally verified seedless binding events. Indeed, those strong-seed targets with seedless MREs were less downregulated by miR-522-3p overexpression than those without seedless MREs (Figure 3F). Taken together, the data above support that the attenuation effect on strong-site-mediated target regulation could be observed using experimentally validated seedless miRNA-binding sites.

**Figure 3.**
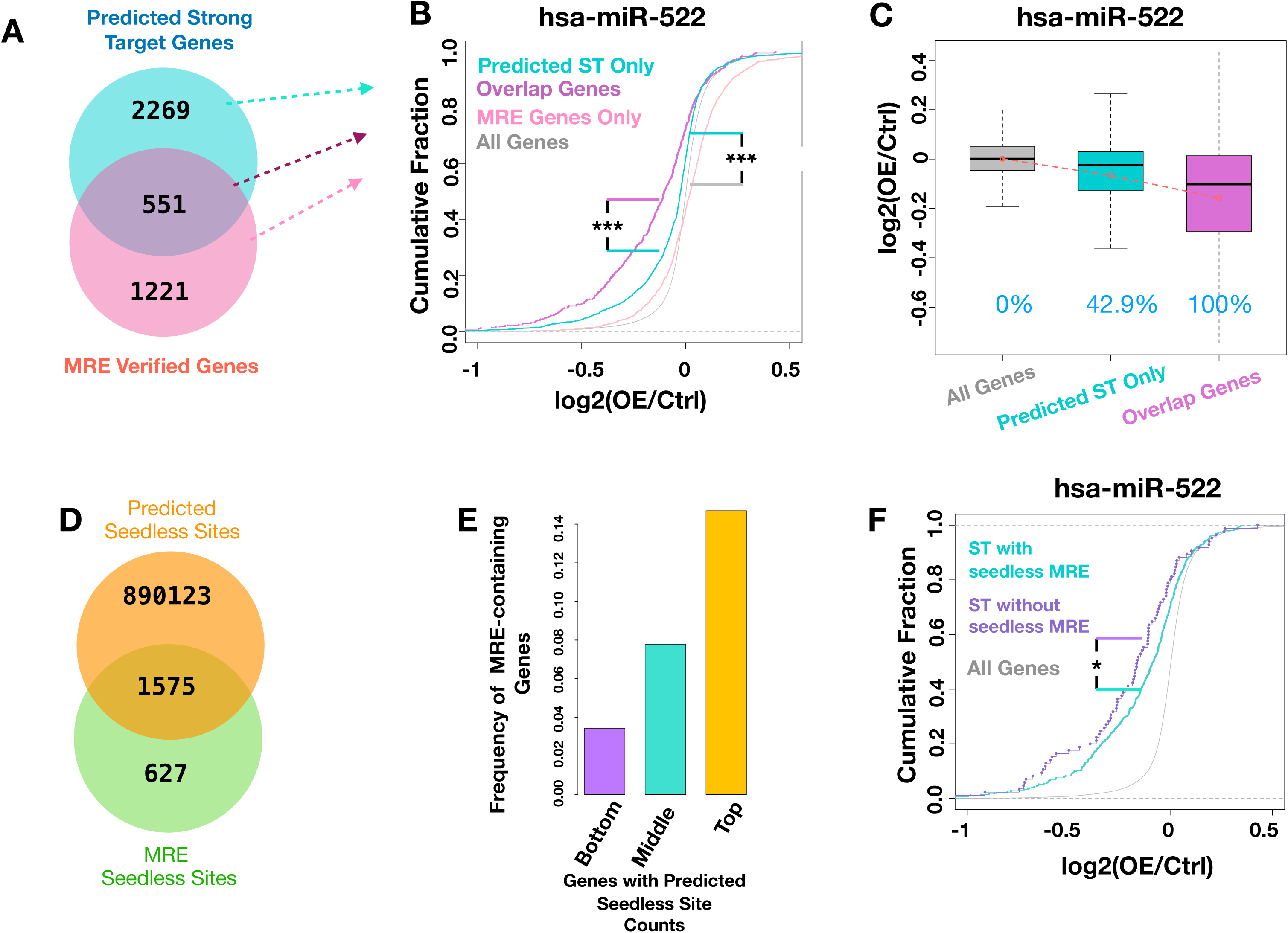
Seedless-site-based attenuation effect can be observed using experimentally verified seedless miRNA binding sites. A dataset was analyzed which documents gene expression and miRNA binding sites (MRE, miRNA responsive elements) after miR-552 overexpression (OE) in MDB-MB-468 cells. **(A)** A Venn diagram showing the relationship between genes containing predicted strong seed sites in the 3’UTRs and MRE-containing genes. **(B)** A cumulative distribution function plot for the indicated groups of genes comparing gene expression in miR-522 OE versus control (Ctrl) cells. ST: strong targets. ***: p<0.001. **(C)** A box and whisker plot for the same data shown in (B), with median levels indicated by the black lines in the boxes and mean level indicated by the dashed-line-connected red dots. Percentage of mean gene expression changes were indicated, with mean levels in all genes set to zero and that in overlap genes set to 100%. **(D)** A Venn diagram showing the relationship between predicted miRNA seedless sites and MRE-verified seedless sites. **(E)** Genes were binned with bottom 1/3, middle 1/3 and top 1/3 of predicted seedless site counts. The observed frequency of MRE-containing genes is plotted which shows a positive correlation with seedless site count. **(F)** A cumulative distribution function plot for miRNA-induced gene expression changes depicting predicted strong targets with or without seedless MRE. *: p<0.05.

To go beyond association results and provide direct causal evidence of seedless-site-mediated attenuation effect, we designed a series of 3′ UTR reporters, in which an 8mer seed site of miR-124 was followed by two copies each of seedless sites for miR-1 and miR-155 (Figure 4A, **Supplementary Table 3**). These reporters were assayed in *DICER1*-null 293T cells in order to reduce the potential influence of seedless interactions by endogenous miRNAs on such reporters. By introducing and varying exogenous miRNA combinations, we would be able to specifically test the impact of seedless sites, without affecting 3′UTR length. We generated nine designs of seedless sites, by varying the sequence complementarity between target site and the miRNA’s non-seed region (Figure 4B-4L). Several seedless designs showed attenuation effects. For example, for design 9, transfecting synthetic miR-124 effectively suppressed reporter activity (Figure 4C, 4D). Synthetic miR-1 did not significantly affect the reporter activity (Figure 4C, 4D), indicating that seedless design 9 is incapable of conferring detectable target suppression by itself under our experimental conditions. However, when miR-124 and miR-1 were co-transfected, there was a complete abolishment of miR-124-induced target suppression (Figure 4C, 4D). This loss of suppression was not due to ineffectiveness of miR-124 when co-transfected with miR-1, because a mutant reporter, in which we mutated the miR-1 seedless sites by altering the sequence corresponding to the non-seed region of the miR-1 (Figure 4B), was suppressed to similar levels under the miR-124/miR-1 co-transfection condition as compared to miR-124 alone (Figure 4C, 4D). To determine if this attenuation effect was due to base-pairing between the non-seed region of miR-1 with the seedless sites, we synthesized a mutant miR-1 whose seed region was identical to miR-1, but the non-seed region was mutated so that it could not match miR-1 seedless sites but could match the mutant miR-1 seedless sites with the same base-pairing pattern consistent with seedless design 9 (Figure 4B). This mutant miR-1 was not able to attenuate miR-124-induced target suppression when co-transfected (Figure 4C, 4D). However, when assayed on the mutant miR-1 seedless reporter, in which the base-pairing in the non-seed region was restored, mutant miR-1 effectively attenuated miR-124-induced target suppression (Figure 4C, 4D). These data support that seedless design 9 could effectively attenuate strong-site-mediated target suppression in a base-pairing dependent manner. We next examined the rest of the seedless designs. Designs 4, 5, 6 and 7 showed similar attenuation effects as design 9 (Figure 4E-4L). Designs 1, 2, 3 and 8, on the other hand, either had weaker or no attenuation effect, or had variable results between experiments (Figure 4E-4L). While detailed rules of functional seedless sites require further exploration, we noticed that the attenuation-competent seedless sites tend to have medium levels of pairing with a wide spectrum of pairing patterns. To demonstrate that the attenuation effect is not restricted to miR-1, we performed similar experiments using miR-155 in combination with miR-124, and observed similar results (**Supplementary Figure S3**). Taken together, the data above support that a wide spectrum of seedless binding patterns could attenuate strong-site mediated target suppression, and further confirm that seedless-site-mediated attenuation is independent from 3′ UTR length.

**Figure 4.**
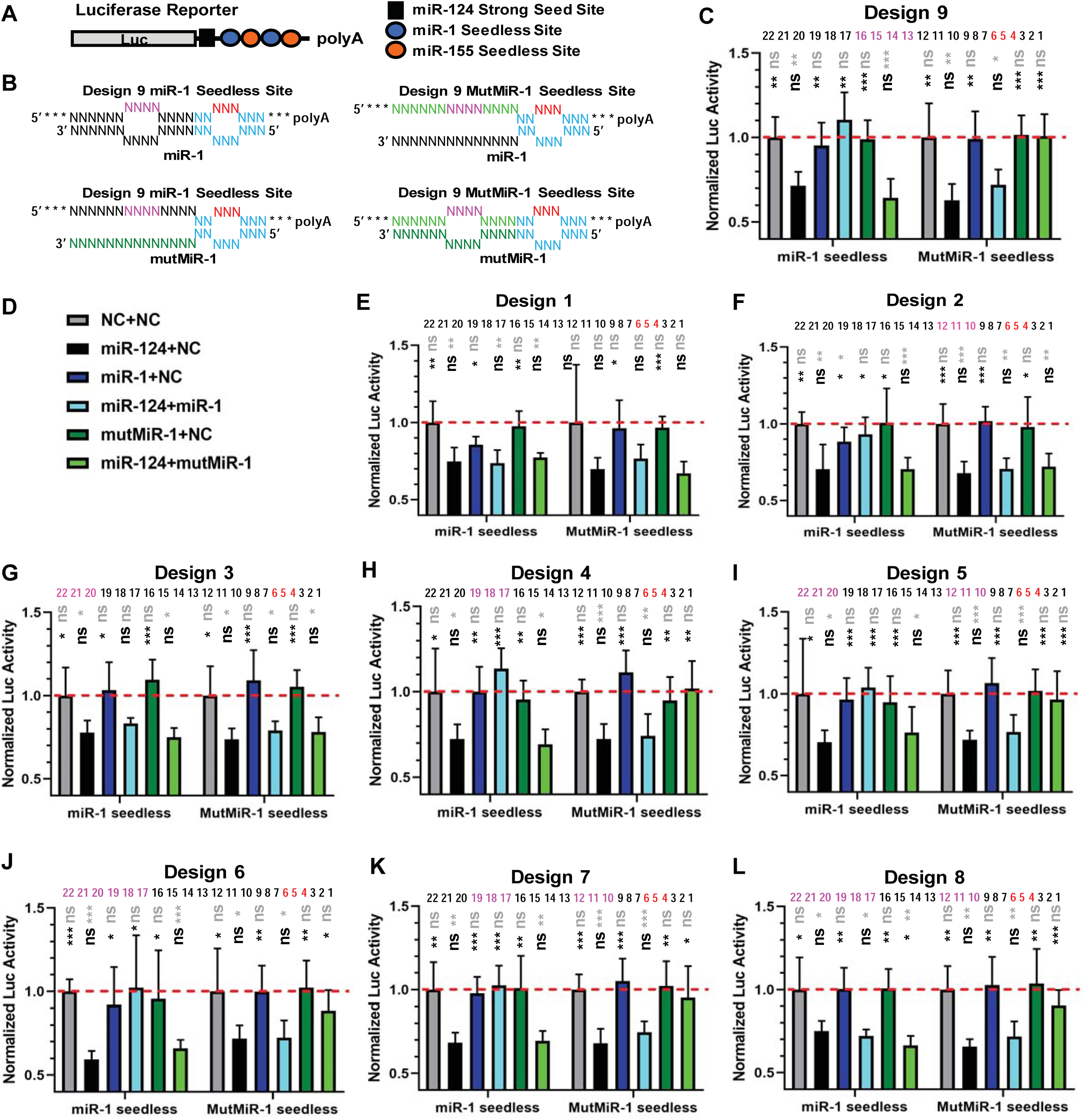
Seedless sites can cause functional attenuation of strong-seed-site-mediated target repression. **(A)** Schematic of the luciferase reporter, with a strong seed site of miR-124 and seedless sites of miR-1 and miR-155 in the 3′ UTR. **(B)** Schematics of pairing patterns of seedless design 9, showing paired and unpaired bases between seedless site and miR-1 sequence, including mutant (mut) seedless site and mutations of miR-1 (mutMiR-1). For miRNA, seed sequence is shown in blue and non-seed region shown in black. Mutated bases are shown in dark green letters. For target site, bases paired with miRNA’s seed region are shown in blue. Red letters indicate that two out of the three red bases are unpaired with miRNA in each of the two copies of the seedless sites. For bases pairing with miRNA’s non-seed region, black letters indicate bases that pair with the wild-type miR-1, magenta letters indicate positions where mutations were designed to disrupt pairing, and light green letters indicate mutant bases that can pair with mutant miR-1. **(C, D)** Luciferase reporter results for seedless design 9. Reporters carrying seedless sites for miR-1 or with mutant seedless sites (MutMiR-1 seedless) were assayed in Dicer-null 293T cells. (D) The legend for the combination of miRNA mimics assayed, with total mimic concentration kept the same across all conditions. NC: negative control mimic. mutMiR-1: mutant miR-1. (C) A position-matching pattern of design 9 is shown above the plot, with seed mismatch bases (two out of the indicated three bases) shown in red and non-seed mismatch bases shown in magenta. Normalized luciferase activities are shown. The red dashed line shows the level of the reporter activity upon treatment of negative control mimic. Statistical significance compared with the grey bar data are shown above the bars in grey font, and those compared with the black bar are shown in black font. *: p<0.05; **: p<0.01; ***: p<0.001; ns: not significant. N=6 except that Design 5 has N=12. Data are representative of two or more experiments. Error bars represent standard deviation. **(E-L)** Similar data as those in (C) are shown for seedless designs 1 to 8, with seedless position-matching patterns indicated.

The reporter data above also indicate that seedless sites of one miRNA could attenuate strong-site-mediated target suppression of another miRNA. If this is the case, we could deduce that when a miRNA is increased in cells, 3′ UTRs without seed site of the miRNA but with high numbers of predicted seedless sites could lead to seemingly miRNA-mediated gene upregulation, with the rationale being that such seedless sites could attenuate endogenous-miRNA-mediated target suppression. Indeed, among the 15 miRNA datasets that showed an attenuation effect, we observed that 8 out of 15 datasets showed a significant association between predicted seedless site count and miRNA-mediated gene expression increase for genes without seed sites in the 3’ UTRs (**Supplementary Figure S4A, Supplementary Table 1**), including two miRNA KD/KO datasets (**Supplementary Figure S4B**). We did notice, however, seedless-site associated gene upregulation for non-seed genes was most prominent when very large numbers of predicted seedless sites are present (**Supplementary Figure S4C**), suggesting potentially a higher functional threshold for one miRNA’s seedless site to attenuate another miRNA’s seed site, as compared to attenuating its own seed site. Among these 15 datasets, only one dataset showed an opposite effect, with larger numbers of predicted seedless sites significantly (p=3.8e-10) associated with miRNA-mediated gene suppression. This observation is consistent with the notion that some seedless sites can suppress gene expression in a context-dependent manner ^14^. Interestingly, among the six datasets that we failed to detect attenuation effects (**Supplementary Table 1**), only one showed an association between seedless site count and gene upregulation of non-seed genes, whereas three out of six showed an opposite association between seedless site numbers and gene suppression. These data suggest that the effect of miRNA-induced gene upregulation is variable and could be partially explained by seedless-site-mediated attenuation effects.

To investigate whether the seedless sites may affect AGO recruitment to strong-site-containing genes, we turned to a published AGO-CLIP dataset in mouse liver, in which AGO-bound miR-122 seed sites have been characterized experimentally through comparison of wildtype and miR-122 knockout liver samples ^35^. Among our computationally predicted miR-122 strong-site-containing targets, 38% had annotated miR-122 binding from the AGO-CLIP data, and these targets were significantly increased upon miR-122 KO (Figure 5A). Similar to the miR-522 MRE dataset, predicted miR-122 strong targets without CLIP-validated binding sites, however, were also significantly upregulated (Figure 5A), again suggesting that a substantial level of false-negatives existed in the AGO-CLIP data. Consistent with the seedless-site-based attenuation of the same miRNA’s seed sites, we observed that among CLIP-validated miR-122 strong target genes, those with higher numbers of predicted seedless sites showed an attenuation effect in target de-repression upon miR-122 KO (Figure 5B). Of note, this dataset was from an independent study from that shown in Figure 1C. In examination of miR-122-induced AGO occupancy signals, as measure by differential read counts between wild-type and miR-122 KO liver among annotated miR-122 seed sites, as expected, we observed that there was a substantially higher AGO occupancy in WT liver than KO liver for predicted strong targets of miR-122 (Figure 5C). However, those 3′ UTRs with more predicted seedless sites showed a significant reduction in AGO occupancy between WT and miR-122 KO liver (Figure 5C), although the effect size was relatively small compared to changes in gene expression (Figure 5B). These data suggest a potential model in which seedless-mediated attenuation reduces AGO occupancy. For 3′ UTRs without predicted miR-122 seed sites, those with lower numbers (bottom 1/3) of predicted miR-122 seedless sites showed no difference in AGO occupancy between WT and KO liver (Figure 5D, 5E). consistent with the notion that AGO occupancy on these 3′ UTRs were predominantly induced by miRNAs other than miR-122. However, for 3′ UTRs without predicted miR-122 seed sites but with higher numbers of predicted miR-122 seedless sites, we observed an increase in AGO occupancy upon miR-122 KO (Figure 5D, 5E), consistent with a model of miR-122-seedless-site-mediated attenuation of AGO occupancy induced by other endogenous miRNAs. The data above suggest that AGO recruitment was reduced with seedless-site-based attenuation.

**Figure 5.**
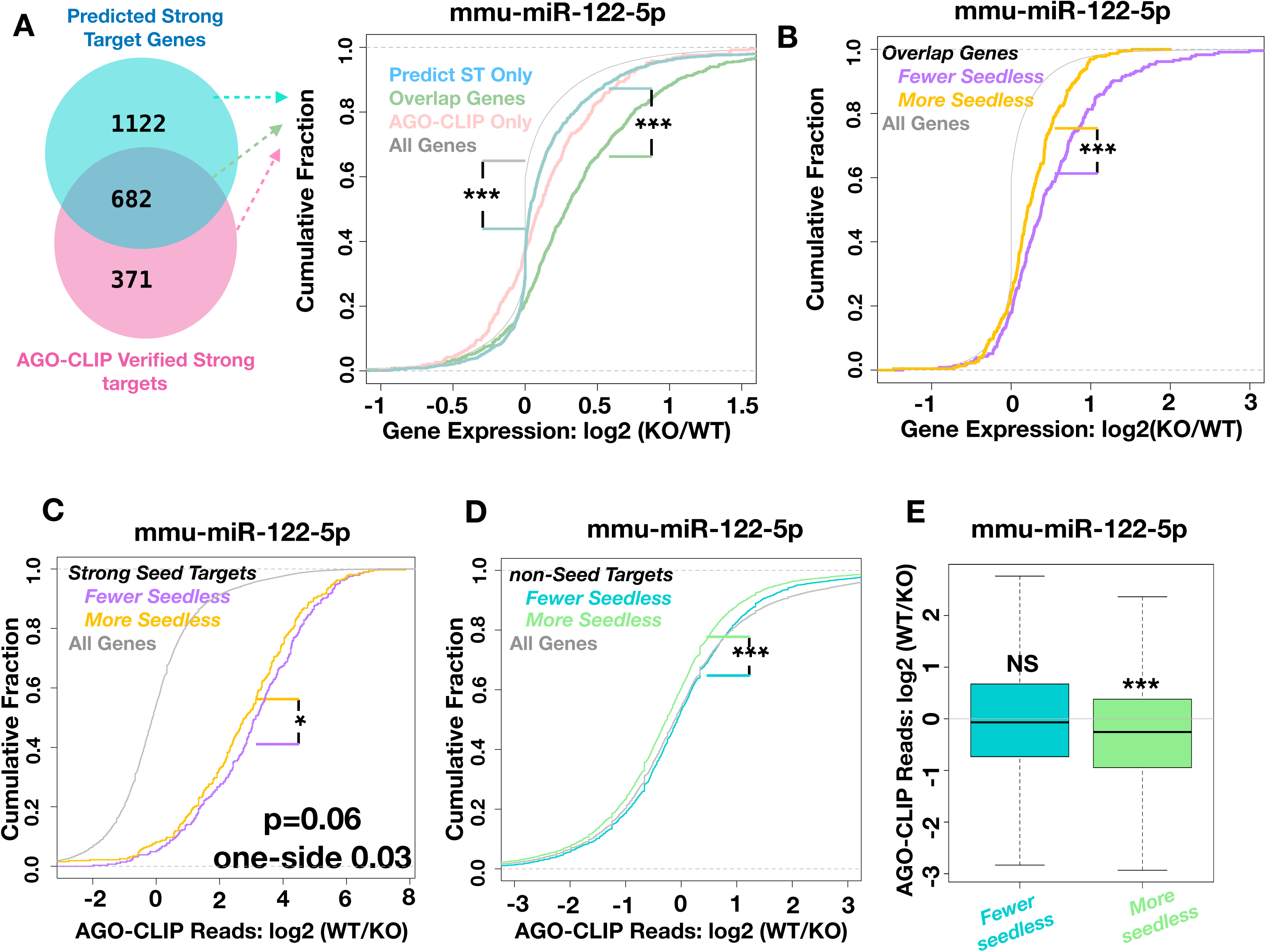
Seedless-site-based attenuation is associated with reduced AGO occupancy on target genes. A dataset was analyzed which documents gene expression and AGO-CLIP data comparing wildtype (WT) and miR-122 knockout (KO) liver. **(A)** Left: A Venn diagram showing the relationship between genes containing predicted miR-122-5p strong seed sites in the 3’UTRs and AGO-CLIP-verified miR-122-5p-bound genes. Right: A cumulative distribution function plot for the indicated groups of genes comparing gene expression in KO versus WT samples. ST: strong targets. ***: p<0.001. **(B)** A cumulative distribution function plot is shown for overlap genes in (A) further divided into those with more or fewer predicted seedless sites (top and bottom 1/3), with all genes as a control. ***: p<0.001. **(C)** Predicted ST genes were separated into those with more or fewer predicted seedless sites, with non-seed genes as a control. The AGO-occupancy difference between WT and KO samples (reflected by difference in log2 of normalized CLIP read counts), is plotted. *: p<0.05. **(D)** A similar analysis as in (C) was carried out for non-seed genes. *: p<0.05. **(E)** The differential AGO-occupancy (KO/WT) for data in (D) was plotted in box and whisker plot, with mean levels indicated.

## Discussion

In this study, we found that miRNA seedless sites in 3′ UTR attenuate strong-seed-site mediated target repression. This attenuation effect could be observed in 15 out of 21 genomic datasets in which the expression of specific miRNAs were increased or reduced, and was independent from 3′ UTR length. We performed reporter assays to show that this attenuation effect by seedless site is causal and dependent on base-pairing between miRNA and seedless target sites. The effect of seedless-site-based attenuation can be strong. In genomic datasets, we observed a difference of ~ two fold or more (Figure 1, **Supplementary Figure S1B, 1D**) between those high-seedless-site-count 3′ UTRs and low-seedless-site-count 3′ UTRs. In reporter assays, seedless sites could completely abolish seed-site-mediated target suppression. It is interesting to note that in our reporter assay for miR-1 seedless sites, the distance between seedless site and seed site falls within the range of synergistic suppression previously observed for two seed sites ^23^, further highlighting the difference in function between seedless sites and seed sites. Occupancy of miRNAs on seedless sites in 3′ UTRs have been frequently observed in biochemical enrichment experiments such as AGO-CLIP or biotinylated miRNA pull down ^4–12^, yet whether most of these biochemically defined miRNA-bound seedless site contribute to gene repression has been questioned recently ^16^. Our model proposes a new function of seedless site that may help to explain the observation of seedless binding by miRNAs. Our model also helps to partially explain miRNA-mediated gene upregulation, and is different from the model of miRNA-miRNA competition for pathway proteins ^36^. We also noted that there were six out of 21 datasets in which we did not see significant seedless-site-based attenuation, suggesting that this attenuation effect may not be applied to all miRNAs or may be dependent on cellular context.

What types of seedless sites can lead to functional attenuation? Our reporter assays using nine seedless designs indicate that there seem to be a wide spectrum of seedless binding patterns that can enable attenuation, yet at the same time not all seedless sites are attenuation-competent. Interestingly, the pairing between miRNA’s 13-16 position is neither required (**Design 9**, Figure 4) nor sufficient (**Design 8**, Figure 4) for functional attenuation, whereas this region has been previously noted to promote miRNA-mediated target suppression ^23^. The computationally predicted seedless sites used in this study likely contain many false positives, yet our analysis of two datasets using biochemical mapping of miRNA binding sites strongly suggests that there are likely many false negatives in biochemically defined miRNA target sites in cells. This notion is not surprising; while these biochemical experiments have been carried out with high technical quality, it is conceivable that weak and transient interactions could be easily missed. Indeed, single molecule experiments have demonstrated that as short as 3 nt matching in the seed sequence is sufficient to trigger efficient recruitment of AGO-miRNA complex to target RNA, despite these shorter matches lead to faster dissociation from target sequence ^37–39^. More sensitive methods are needed to better define weak and transient interactions between miRNAs and target mRNAs globally. In the absence of such a technology, the use of our computationally predicted seedless sites can be viewed as a surrogate reflecting the likelihood of seedless interactions rather than an absolute indication of the existence of seedless interactions. Further characterization of the rules underlying functional seedless sites in the future could also help to generate better computational prediction for seedless-site-based attenuation.

There are a number of potential mechanisms by which seedless sites may affect strong-site-mediated target repression. For example, it is possible that seedless sites may serve as transient decoys, possibly serving as bind-and-leave sinks during the lateral diffusion of miRNAs ^37,38,40^ on target mRNA, thus preventing miRNA from finding its seed sites. This possibility is consistent with our observation of reduced AGO binding in the presence of large numbers of predicted seedless sites. Another possibility is that the effect of seedless sites is not mediated solely through AGO. This possibility is suggested by the lack of requirement of pairing involving miRNA’s 13-16 position for functional attenuation. Indeed, several studies have proposed the existence of a pool of AGO-free miRNAs inside cells ^32–34^. It is thus conceivable that AGO-free miRNAs may recruit additional trans-factors to impede strong-site-mediated AGO occupancy, leading to reduced target suppression. The detailed mechanism of seedless-site-mediated attenuation will require further investigation.

## Methods

### Constructs

All luciferase reporters were cloned into the psiCHECK2 vector (Promega, #C8021) using NotI and XhoI restriction enzyme sites. Sequences that are cloned between these two sites are shown in **Supplementary Table 3**.

### Luciferase Reporter Assay

Luciferase reporter assays were carried out in Dicer-KO 293T cells ^41^ (NoDice 4-25 clone, kind gift from Dr. Bryan Cullen). Cells were cultured in DMEM medium (Gibco, #10569010) with 10% FBS (Gibco, #16140071) and 1% Penicillin-Streptomycin-Glutamine (Gibco, #10378016).

The transfection was carried out by either one-step or two-step procedures. For one step transfection, cells were plated in 384-well plate wells, at 3000 cells per well. On the second day, cells in each well were co-transfected with 6 ng of reporter plasmid,100 nM of a mixture of miRNA mimics, and 0.038ul Lipofectamine 3000 (Invitrogen, #L3000015) following manufacturer’s protocol. For each miRNA, the final concentration is 50 nM for each assay, and when necessary, 50 nM Negative Control mimic was added to maintain the same final concentration of small RNAs (100 nM) per well. Two days after the small RNA transfection, luciferase assays for both firefly and renilla luciferase activity were performed using the Dual-Glo kit (Promega, #E2940) following the manufacturer’s protocol. For two-step transfection, cells were plated in 12-well plate wells, at 5×10^5^ cells per well. On the second day, cells were transfected with reporter plasmids, with each well transfected with 100 ng of plasmid and 1.5 µl Lipofectamine 3000 following manufacturer’s protocol. After an overnight culture, cells from each 12-well-plate well were trypsinized and replated at 3000 cells/well in 384-well plates. On the next day, 100 nM of miRNA mimic mixture was transfected into each well. After two days, luciferase assays were performed using Dual-Glo Luciferase kit.

Double stranded miRNA mimics were purchased from Dharmacon. These mimics were synthesized using ON-Target modification for strand-specific AGO loading. Wildtype hsa-miR-124-3p, hsa-miR-1-3p and hsa-miR-155-5p were purchased from catalog, whereas mutMiR-1 was custom synthesized with the following ON-Target sequence: UGGAAUGUUGUAGUAAGAAAUA. The design of mutMiR-1 had the first 8 bases the same as in miR-1, but the non-seed region was scrambled so that the overall A, U, C and G content remained the same as in miR-1, but every base in the non-seed region was mutated. Control mimic was Negative Control #1 (Invitrogen, #4464058) which was a double-stranded miRNA mimic with random sequence.

Three types of control assays were built into each 384-well plate, including CtrlUTR-CtrlMiR assays (a control reporter assayed with a negative control miRNA mimic), UTR-CtrlMiR assays (experimental reporters assayed with negative control miRNA mimic), and CtrlUTR-miR assays (a control reporter assayed with each of the miRNA mimics on the plate). Control reporter was psiCHECK2 without cloned 3′ UTR.

We then compute the ratio of renilla luciferase versus firefly luciferase readings (RvF ratios). The RvF ratio of any given well, including controls, was then normalized using the following formula. Normalized Luciferase Activity = (RvF_Well_ / mean(RvF_UTR-CtrlMiR_))/(mean(RvF_CtrlUTR-miR_)/mean(RvF_CtrlUTR-CtrlMiR_)). After normalization, the means of all three control assays became one.

### Predictions of miRNA binding sites

For miRNAs in this study, the STarMir program ^42^ was used to make predictions of both seed and seedless binding sites on mRNAs. The program was based on modeling of data from CLIP studies ^18^, and incorporated the RNAhybrid program ^43^. All possible binding sites from RNAhybrid that have perfect match to the miRNA’s seed region were included except for those with GU base pairing that violates the seed rule. Predicted seedless binding sites with hybridization energy of ≤ −15 kcal /mol were included in our analyses, as the number of predicted seedless sites was large and the threshold of −15 kcal /mol for a stable hybrid was previously established^17^. Seed sites were further classified into 8mer, 7mer-m8, 7mer-A1, 6mer and offset-6mer based on the predicted binding patterns ^3^. Although a probability of a predicted site being a miRNA binding site is available from STarMir, we did not use this probability to filter the predicted sites.

In this study, we define strong seed sites as those predicted sites that belong to 8mer, 7mer-A1 or 7mer-m8 categories. We define strong-seed-targets as those genes whose 3′ UTR harbors at least one strong seed site for a given miRNA. We define non-seed genes as those genes whose 3′ UTR does not have any type of predicted seed sites for a given miRNA.

### Data Sources and Data Processing

MiRNA knockdown or over-expression data were downloaded from GEO database, which included GSE16568, GSE16700, GSE16569, GSE92564, GSE86575, GSE31397, GSE85884, GSE85884, GSE7333, GSE85884, GSE28456, GSE92564, GSE79340, GSE16572, GSE85884, GSE39779, GSE16571. For data from the Affymetrix microarrays, raw data were downloaded and normalized by the RMA method using R package “affy”. Custom CDF files from Brainarray database were used to map the probes into RefSeq transcripts (http://brainarray.mbni.med.umich.edu/Brainarray/Database/). For data from other platforms. Processed data were directly download by the R “GEOquery” package.

Data related to miR-124, miR-122, miR-522 and miR-223, as well as the processed AGO-CLIP data and processed MREs IMPACT-seq data were obtained from relevant supporting tables published in the original papers (see **Table S1**). Annotation of the binding sites for miR-122 was directly obtained from tables in the published study. For all these datasets, gene IDs were converted into RefSeq ID to cope with the prediction results by the R package “biomaRt”.

For annotating the miR-522 binding sites, MRE sequences and genomic coordinates (hg19) were obtained from the publication ^4^. We then mapped our predicted target site sequences to the reference genome (hg19) by the STAR software to obtain their genomic locations. The intersection of experimental MRE regions of miR-522 and predicted sites were calculated by intersectBED function from BEDtools. Given the size of MREs, the predicted sites located within a genomic distance of 100bp to MRE were defined as overlaps.

For the luciferase reporter data on TET2 and Tet2 3′ UTRs, the effects of miRNA overexpression on luciferase reporter were downloaded from the original study ^31^, and the mean normalized luciferase activity was used.

### Analysis of the Effect of Seedless Sites and 3’UTR Length Using Cumulative Distribution Functions

Cumulative distribution function (CDF) plots were used to visualize the gene expression difference between groups of genes upon OE or KO of a given miRNA. For miRNA OE or KD datasets, log2 fold change of OE samples over control samples were used. For miRNA KO datasets, log2 fold change of KO samples over WT samples were used. If there were multiple samples that below to KO or WT groups, mean expression was calculated for KO or WT groups for each gene before further analysis. For analysis associated with evaluating one miRNA’s seedless sites on the same miRNA’s seed site, analysis was restricted to those genes whose 3’UTR harbors at least one strong seed site for a given miRNA, before subcategorizing genes based on whether they have seedless site count in the top one-third or bottom one third of counts. For analysis evaluating miRNA-induced gene upregulation, analysis was restricted to non-seed genes. For 3′ UTR length analysis, similar approaches were taken based on 3′ UTR length.

For the luciferase reporter data on TET2 and Tet2 3′ UTRs, we categorized miRNAs based on whether there are predicted strong seed sites and the number of predicted seedless sites. For analysis of the effect of one miRNA’s seedless site on its own seed site, we filtered miRNAs to retain those that at least one predicted strong seed site in the 3′ UTR. These miRNAs were further categorized based on the number of seedless sites by ranking them according to the number of seedless sites. Those miRNAs with the top one-third of seedless site counts were compared to those with the bottom one-third of seedless site counts. For analysis of the effect of miRNA’s seedless site on gene upregulation, we filtered miRNAs to retain those that do not have any type of seed sites in the 3′ UTR, before further categorizing them based on the number of seedless sites.

The R function “ecdf()” was utilized to generate the plots, with p-value calculation detailed in the Statistics section.

### Linear Regression Modeling

Linear regression analyses were conducted to assess the contribution of the number of predicted miRNA seedless sites or 3′ UTR length to miRNA-mediated gene regulation. For miRNA OE or KD datasets, fold changes were assessed between OE samples and controls samples. For miRNA KO datasets, fold changes were assessed between KO samples and WT samples.

In the univariate model setting, the model was fitted as follows:

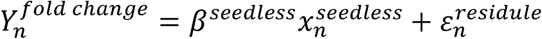

In the above formula, 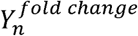 is the log2 fold change for gene n. 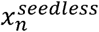 is the count of the predicted seedless sites for the given miRNA in the 3′ UTR of gene n. 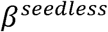 is the weight to be fitted. 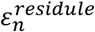 is the random error term.

The effects of seedless sites on gene expression fold changes were assessed by the significance of *β*^*seedless*^.

When evaluating the attenuation effects by one miRNA’s seedless sites on the same miRNA’s strong seed sites, we limited the collection of genes for model fitting to those that have predicted strong seed sites in the 3′ UTR.

When evaluating the miRNA-mediated upregulation effects, we limit the collection of genes for model fitting to that do not have any type of predicted seed sites in the 3′ UTR.

When evaluating the contribution of 3′ UTR length, a similar formula was used, with corresponding terms for 3′ UTR length replacing those for seedless site count.

For assessing the effect of seedless sites or 3′ UTR length separately in the meta-data analysis, the following model was used:

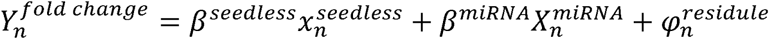

In the above formula, 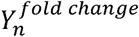 is the log2 fold change for gene n. 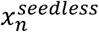 is the seedless site count for gene n. 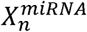 is a n*(m-1) matrix of dummy variables indicating whether the data point was from a dataset in which a particular miRNA was perturbed; n is the count of genes and m is the count of miRNA datasets. The effect of seedless sites was evaluated through the significance of *β*^*seedless*^. For assessing the effect of 3′ UTR length, a similar formula was used by replacing seedless terms with those for 3′ UTR length.

For assessing the effects of both seedless sites and 3′ UTR length simultaneously in the meta-data analysis, the following model was used:

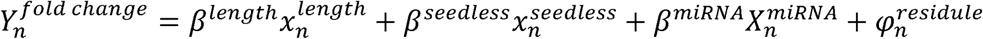

In the above formula, 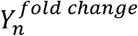 is the log2 fold change for gene n. 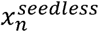 and 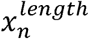 are two variables representing the seedless site counts and 3′ UTR length, respectively. 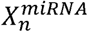 is a n*(m-1) matrix of dummy variables indicating whether the data point was from a dataset in which a particular miRNA was perturbed; n is the count of genes and m is the count of miRNA datasets. The effect of seedless sites was evaluated through the significance of *β*^*seedless*^ by accounting for the effect of 3′ UTR length as well as the baselines of various miRNAs.

Data fitting was performed by using the R function “glm()”, with the co-efficiencies of the model calculated by minimizing the sum of squared errors.

### Statistics

To evaluate whether one CDF distribution is significantly left-shifted or right-shifted in comparison to another, we performed one-sided two sample KS test, using the R function “ks.test()”. For example, for evaluating the attenuation effects by the seedless sites, we tested whether the genes with more seedless sites had significantly reduced strong-seed-site-mediated suppression in comparison to those genes with fewer seedless sites. For evaluation of significance of linear regression, p-values were obtained from the R function “glm()”. For Figure 5E evaluating changes in AGO occupancy upon miR-122 KO dataset, one sample t-test was used.

For calculating significance of the luciferase reporter data, two sample student’s t-test was performed using Microsoft Excel, with a two-sided test assuming unequal variance.

## Acknowledgement

We thank William Rennie and Shaveta Kanoria for assisting with preparation of miRNA binding site prediction data. Services provided by the NIDDK-supported Yale Cooperative Center of Excellence in Hematology (U54DK106857) assisted this study.

## Author Contributions

JL, HL and HZ supervised the study. YD supervised provision of prediction data of miRNA target sites, prepared the description of the prediction procedure, and reviewed the manuscript. SM and JL made the initial observation of the association between seedless sites and strong-seed-site-mediated gene regulation. XW and PC performed all bioinformatics analyses, assisted by DZ and JT. JT performed all bench experiments, with XW, PC, DZ and JL participating in experimental design. JL, XW, PC and TJ wrote the manuscript.

