## Supplemental Figures for "microRNA seedless sites attenuate strong-seed-site-mediated target repression"

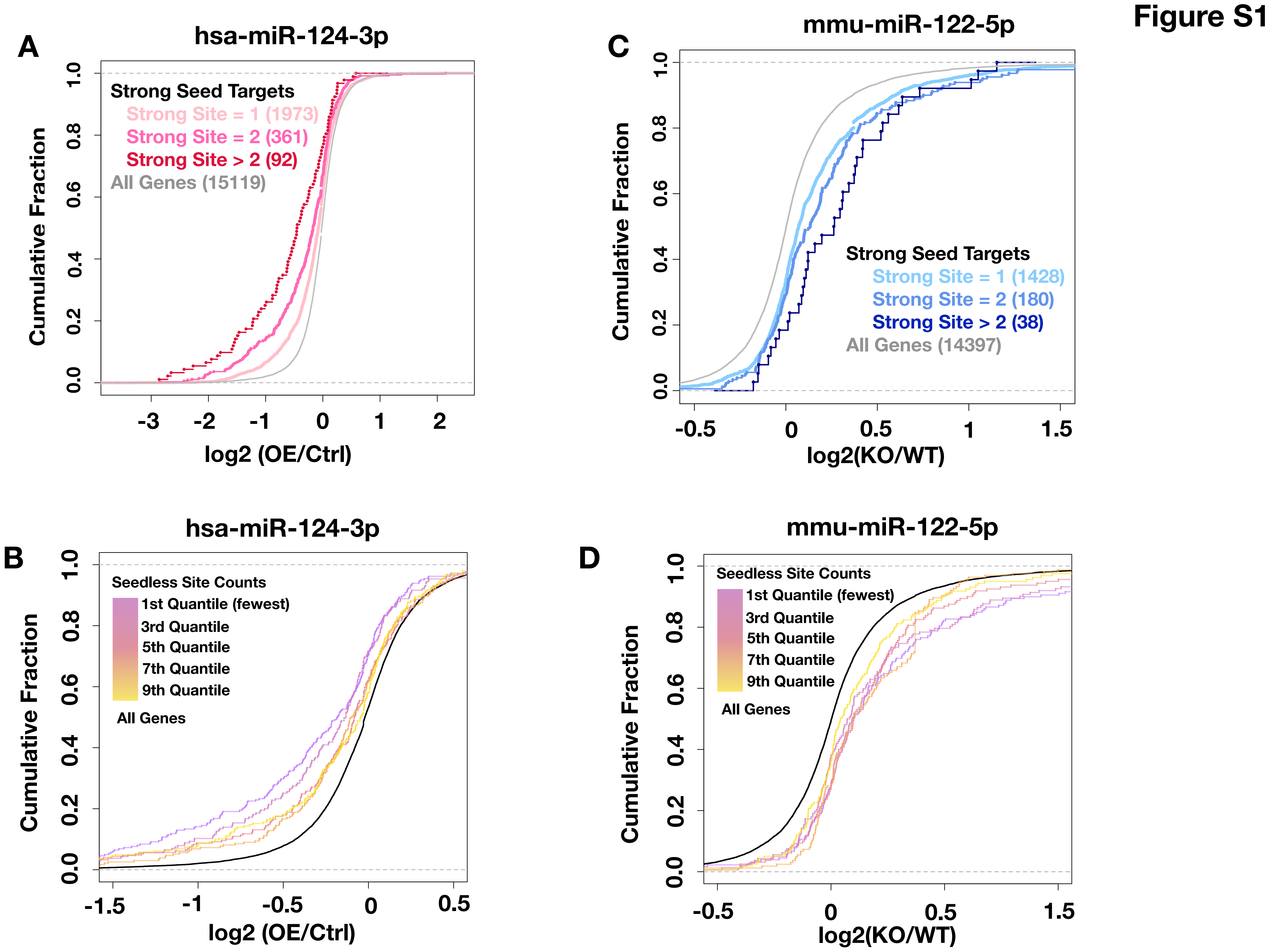
Figure S1
A
hsa-miR-124-3p
Strong Seed Targets
Strong Site = 1 (1973)
Strong Site = 2 (361)
Strong Site > 2 (92)
All Genes (15119)
Cumulative Fraction
log2 (OE/Ctrl)
C
mmu-miR-122-5p
Strong Seed Targets
Strong Site = 1 (1428)
Strong Site = 2 (180)
Strong Site > 2 (38)
All Genes (14397)
Cumulative Fraction
log2(KO/WT)
B
hsa-miR-124-3p
Seedless Site Counts
1st Quantile (fewest)
3rd Quantile
5th Quantile
7th Quantile
9th Quantile
All Genes
Cumulative Fraction
log2 (OE/Ctrl)
D
mmu-miR-122-5p
Seedless Site Counts
1st Quantile (fewest)
3rd Quantile
5th Quantile
7th Quantile
9th Quantile
All Genes
Cumulative Fraction
log2(KO/WT)

### Figure S1 Continued

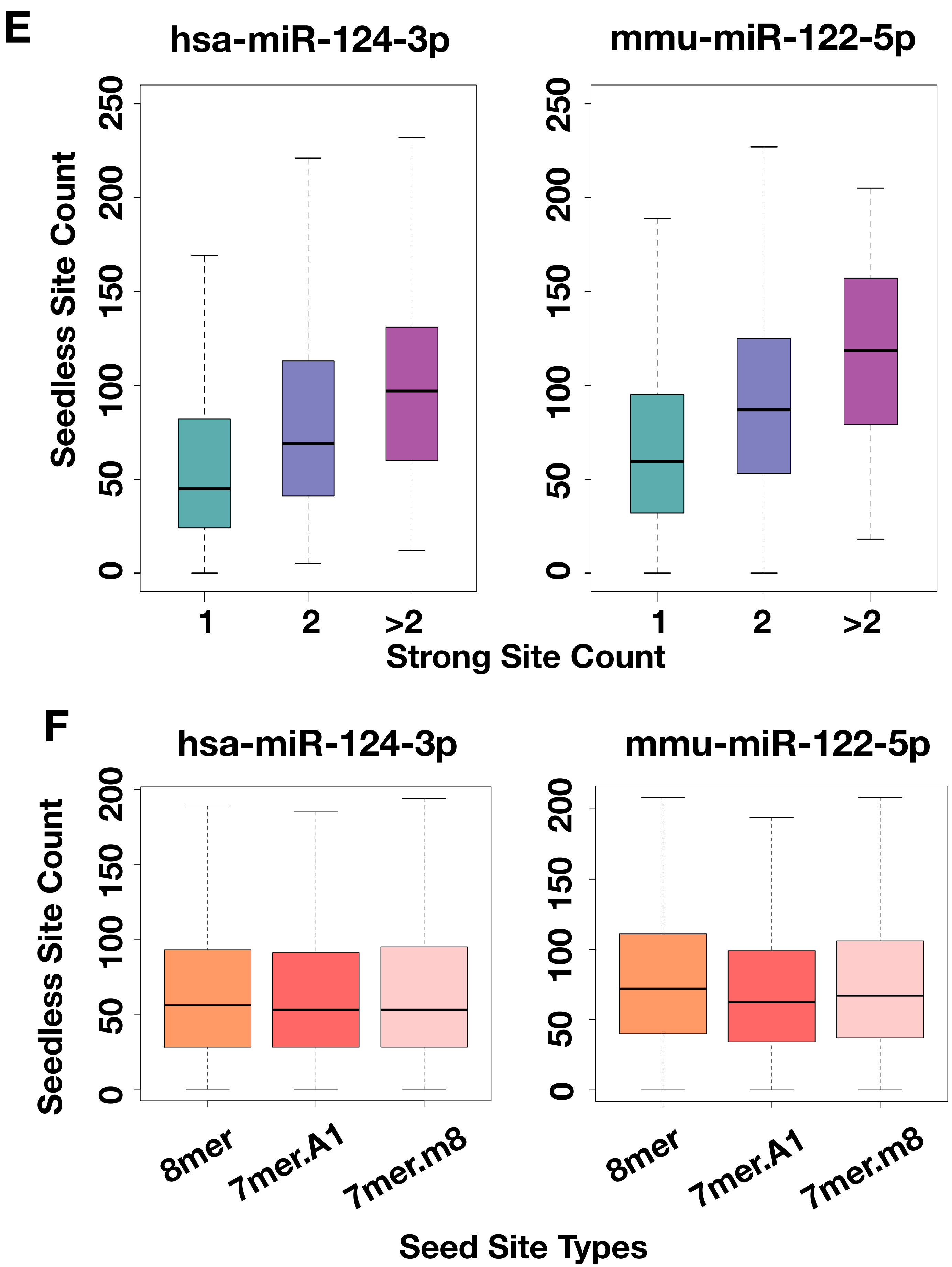

**Figure S1. The impact of seed and seedless sites on miRNA-induced gene expression changes. (A)** Predicted miR-124-3p strong seed targets were further separated into those containing 1, 2 or more strong seed sites. A CDF plot is shown depicting gene expression changes for the indicated groups after miR-124 overexpression in HeLa cells. Number in parentheses indicates the number of genes in the corresponding group. **(B)** Predicted strong targets of miR-124-3p were further separated into 10 bins with regard to predicted seedless site count. A CDF plot shows a graded response of seedless site effect, with only odd numbered bins shown to avoid clutter. **(C, D)** Similar plot as in (A) and (B), but plotted for miR-122 knockout (KO) liver versus wildtype (WT) samples. **(E)** Strong-seed targets of the indicated miRNA were separated into bins according to the number of predicted strong seed sites. The numbers of predicted seedless sites in these bins are shown in a box and whisker plot, revealing a positive correlation between seedless site count and predicted seed site count. This association cannot explain the observed attenuation effects. **(F)** Strong-seed targets of the indicated miRNA, with a single strong seed in the 3'UTR, were separated according to seed site types. The predicted seedless site counts for these groups are plotted in a box and whisker plot. No strong association was observed.

### Figure S2

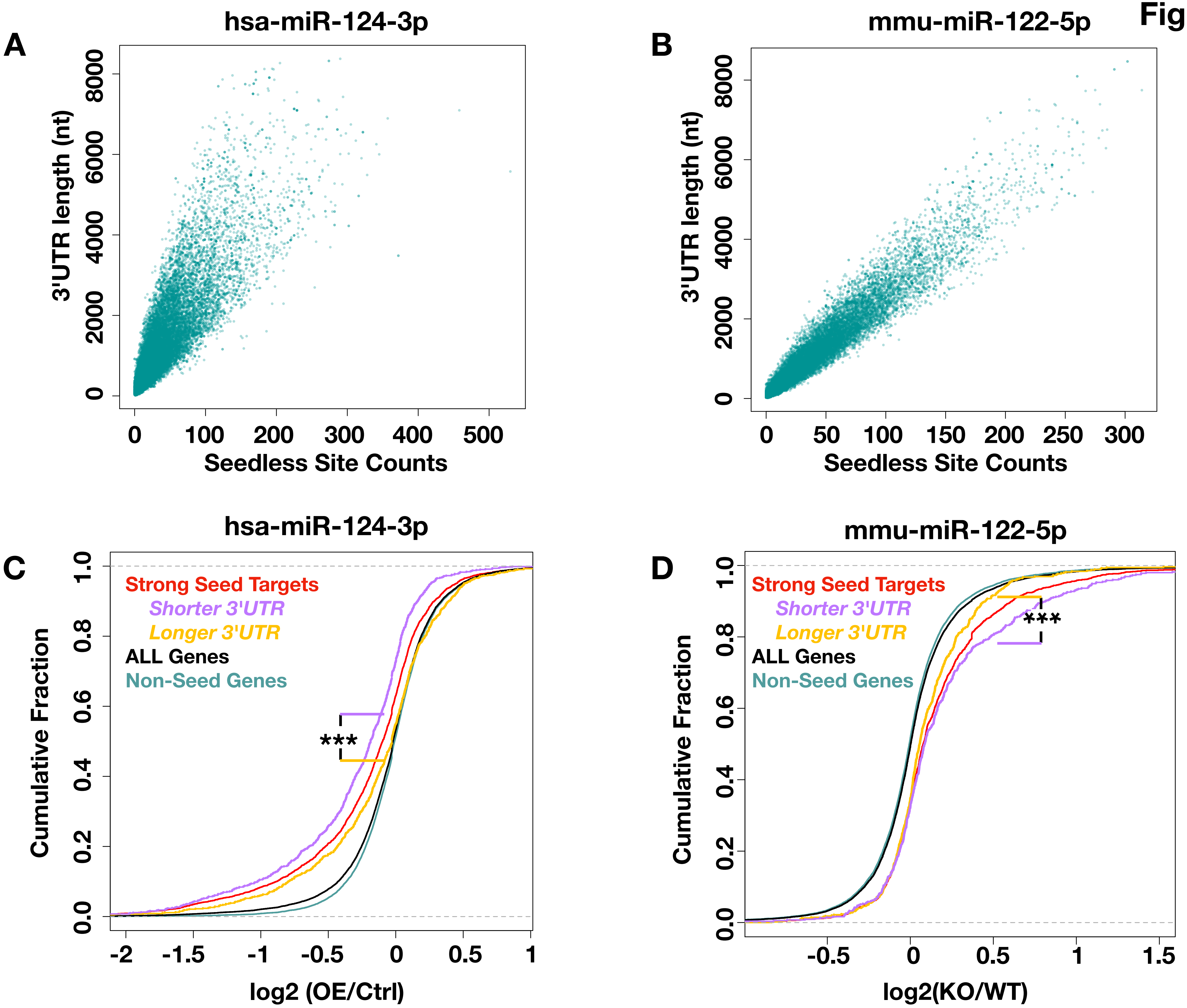

A
hsa-miR-124-3p
3'UTR length (nt)
0
2000
4000
6000
8000
0
100
200
300
400
500
Seedless Site Counts
B
mmu-miR-122-5p
3'UTR length (nt)
0
2000
4000
6000
8000
0
50
100
150
200
250
300
Seedless Site Counts
C
hsa-miR-124-3p
Cumulative Fraction
0.0
0.2
0.4
0.6
0.8
1.0
Strong Seed Targets
Shorter 3'UTR
Longer 3'UTR
ALL Genes
Non-Seed Genes
***
-2
-1.5
-1
-0.5
0
0.5
1
log2 (OE/Ctrl)
D
mmu-miR-122-5p
Cumulative Fraction
0.0
0.2
0.4
0.6
0.8
1.0
Strong Seed Targets
Shorter 3'UTR
Longer 3'UTR
ALL Genes
Non-Seed Genes
***
-0.5
0
0.5
1
1.5
log2(KO/WT)

**Figure S2. The effect of 3'UTR length on miRNA-induced gene expression changes. (A, B)** Dot plots are shown for the relationship between the number of predicted seedless sites and 3'UTR length, with each dot representing a gene, for (A) hsa-miR-124-3p and (B) mmu-miR-122-5p. 3'UTR length is measured in nucleotides (nt). **(C, D)** Predicted strong seed targets were further divided into those with longer and shorter 3'UTRs (top and bottom 1/3 of genes, respectively), and compared for miRNA-induced gene expression changes. Cumulative distribution function plots for the indicated groups of genes comparing gene expression in (C) miR-124 overexpression (OE) versus control (Ctrl) HeLa cells, and (D) miR-122 knockout (KO) liver versus wildtype (WT) liver. Non-seed genes refer to genes without predicted seed sites in the 3'UTR. Legends are color-coded to match the line colors. ***: $p<0.001$.

### Figure S3

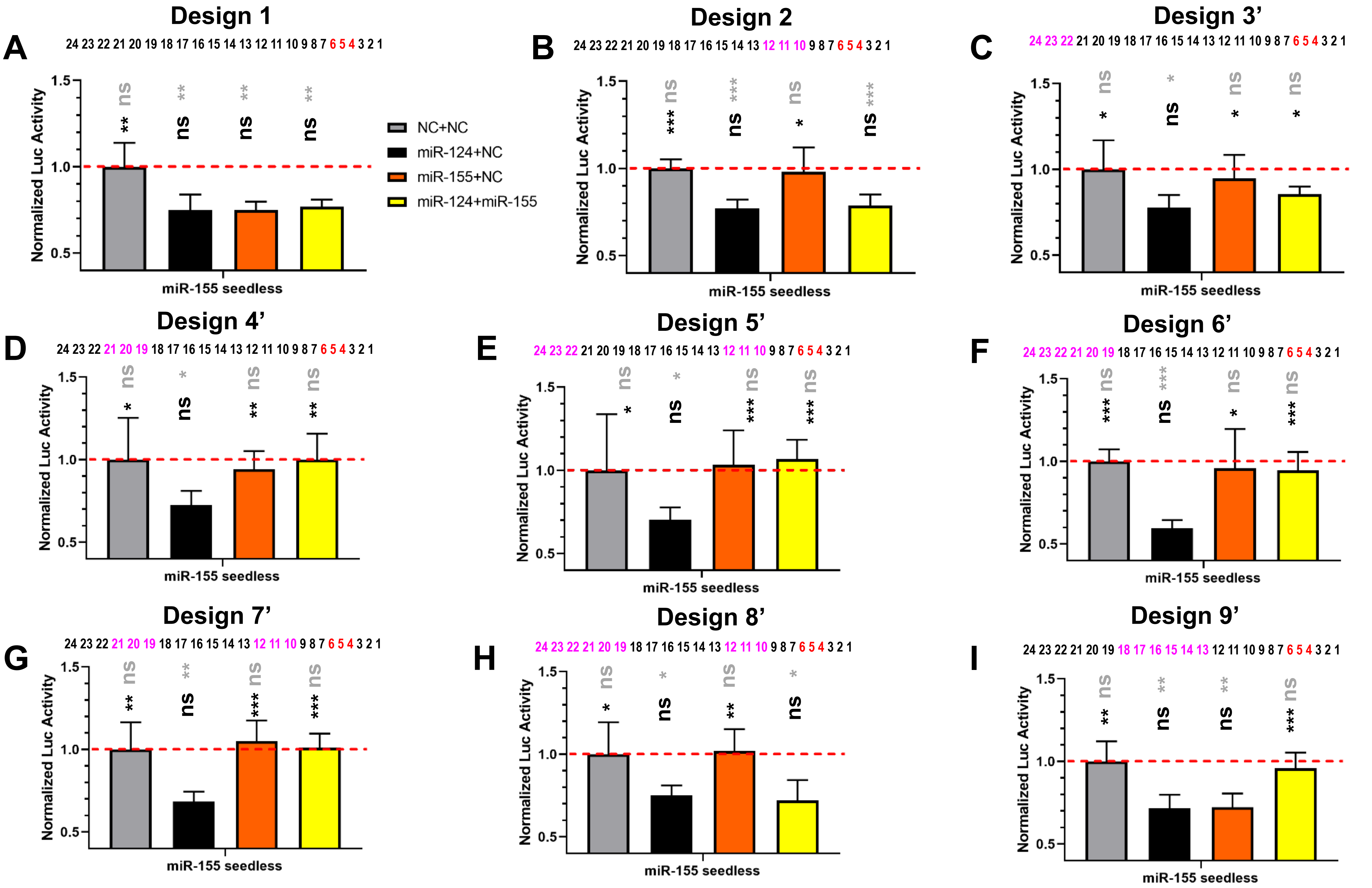
A
B
C
D
E
F
G
H
I
Design 1
Design 2
Design 3'
Design 4'
Design 5'
Design 6'
Design 7'
Design 8'
Design 9'
24 23 22 21 20 19 18 17 16 15 14 13 12 11 10 9 8 7 6 5 4 3 2 1
Normalized Luc Activity
miR-155 seedless
NC+NC
miR-124+NC
miR-155+NC
miR-124+miR-155
ns
*
**
***

**Figure S3. Experimental validation of seedless-site-mediated attenuation. (A)** Luciferase reporter results for seedless design 1. Reporters carrying seedless sites for miR-155 were assayed in Dicer-null 293T cells. The legend for the combination of miRNA mimics assayed is shown to the right, with total mimic concentration kept the same across all conditions. NC: negative control mimic. A position-matching pattern of design 1 is shown, with seed mismatch bases (two out of the indicated three bases) shown in red and non-seed mismatch bases shown in magenta. Normalized luciferase activities are shown. The red dashed line shows the level of the reporter activity upon treatment of negative control mimic. Significance levels compared with the grey bar data are shown in grey font, and those compared with the black bar are shown in black font. *: $p<0.05$; **: $p<0.01$; ***: $p<0.001$; ns: not significant. N=6 except that Design 5 has N=12. Error bars represent standard deviation. **(B-I)** Similar data as those in (A) are shown for seedless designs 2 and 3' to 9', with position-matching patterns indicated.

**Figure S4**

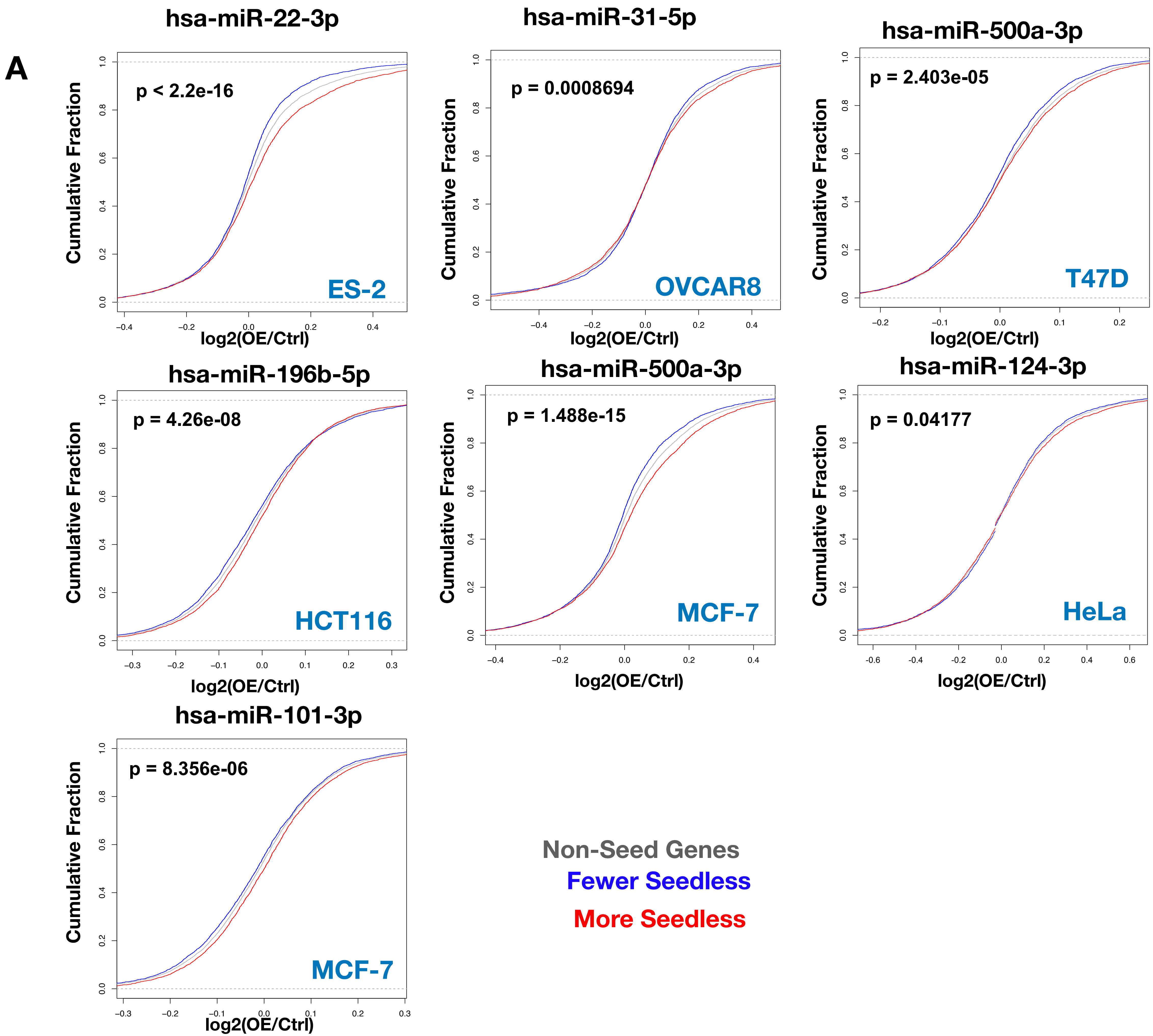
A
hsa-miR-22-3p
p < 2.2e-16
ES-2
hsa-miR-31-5p
p = 0.0008694
OVCAR8
hsa-miR-500a-3p
p = 2.403e-05
T47D
hsa-miR-196b-5p
p = 4.26e-08
HCT116
hsa-miR-500a-3p
p = 1.488e-15
MCF-7
hsa-miR-124-3p
p = 0.04177
HeLa
hsa-miR-101-3p
p = 8.356e-06
MCF-7
Cumulative Fraction
log2(OE/Ctrl)
Non-Seed Genes
Fewer Seedless
More Seedless

### Figure S4 Continued

B

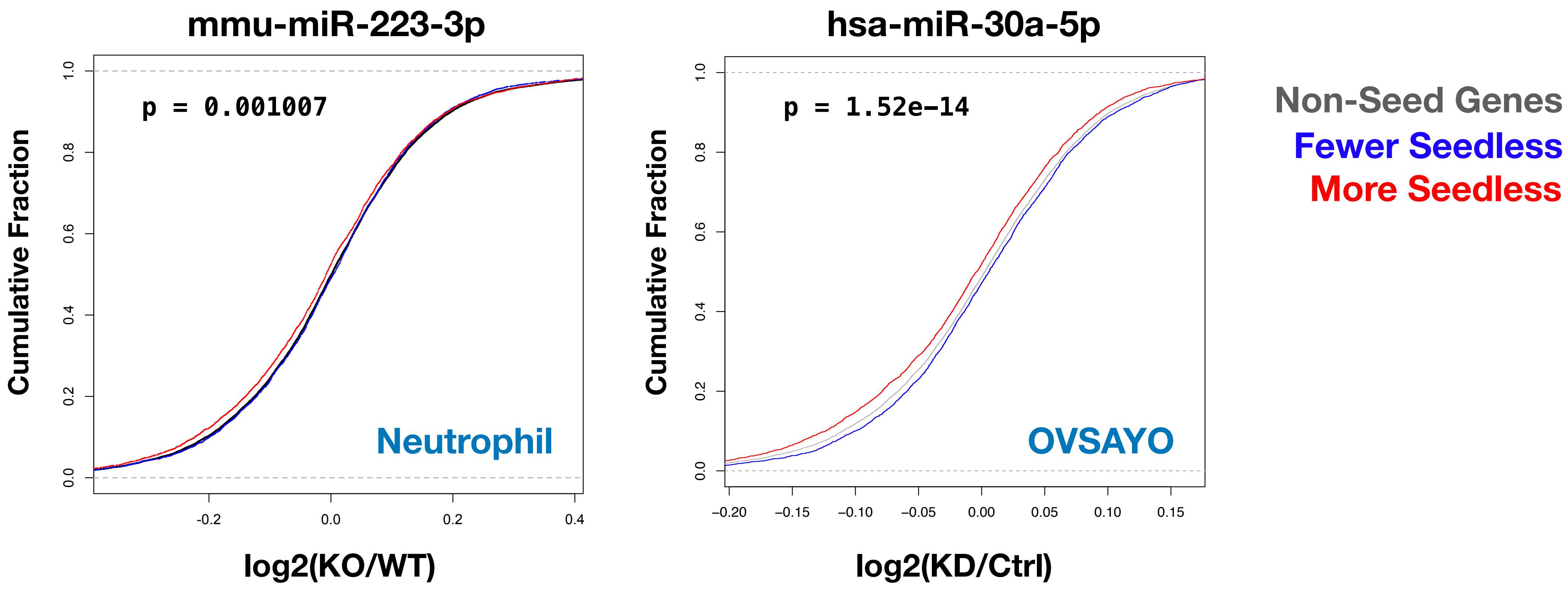


C

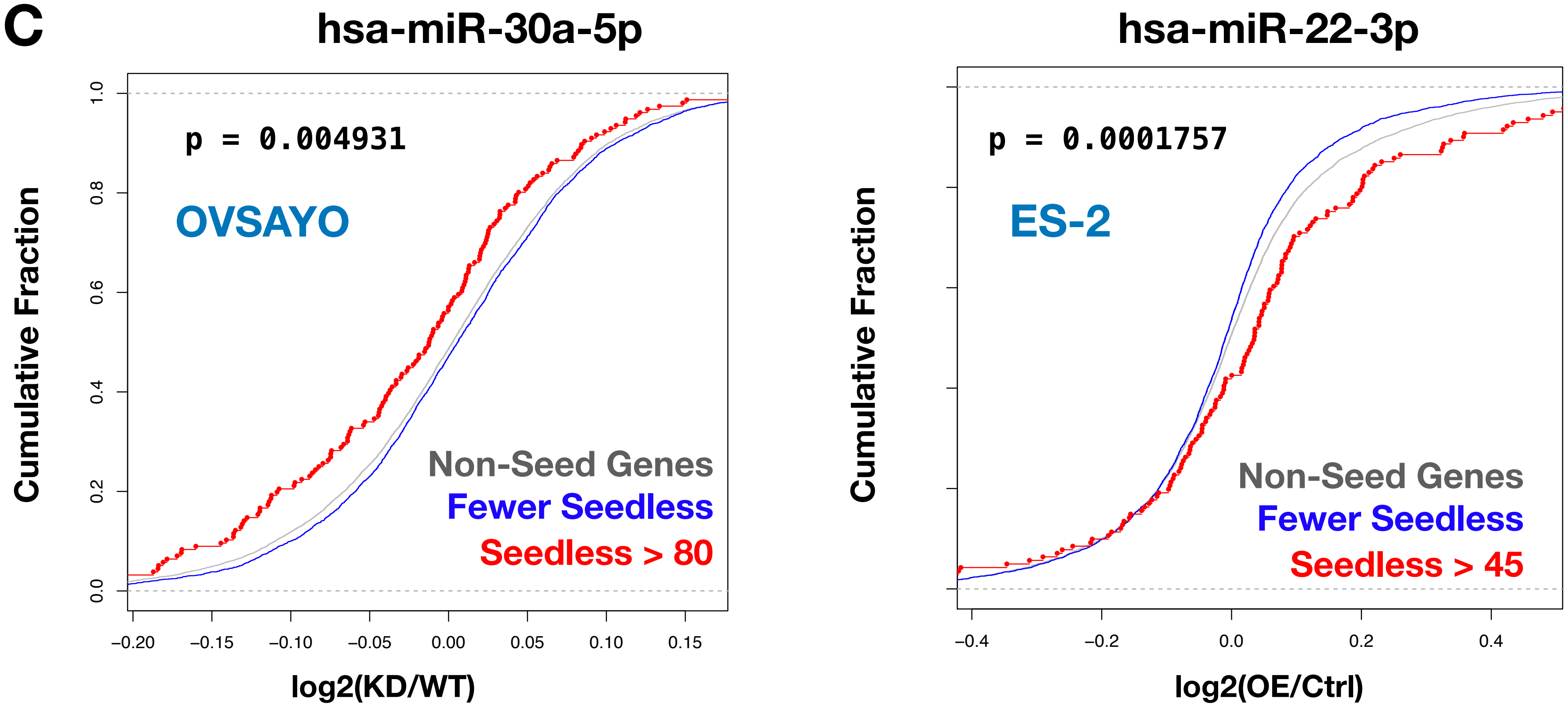

**Figure S4. Higher seedless site count is associated with miRNA-mediated gene upregulation of non-seed genes. (A)** Panels show CDF plots for miRNA overexpression (OE) datasets that had significant association between seedless site count and miRNA-mediated gene upregulation of non-seed genes. Genes without predicted seed sites in the 3'UTR for the indicated miRNA (non-seed genes) were further separated into those that have more or fewer predicted seedless sites for that miRNA (top and bottom 1/3 of genes). Line legend is shown at the bottom right corner. miRNA name, cell type name and p values (between red and blue lines) are indicated in each panel. **(B)** Similar to (A), miRNA knockout (KO) or knockdown (KD) data are shown. **(C)** Two examples are shown in which very large seedless numbers are associated with more obvious gene upregulation of non-seed genes.
